# GalDrive: Pipeline for comparative identification of driver mutations using the Galaxy framework

**DOI:** 10.1101/010538

**Authors:** Saket K Choudhary, Santosh B Noronha

## Abstract

Identification of driver mutations can lead to a better understanding of the molecular mechanisms associated with cancer. This can be a first step towards developing diagnostic and prognostic markers. Various driver mutation prediction tools rely on different algorithm for prediction and hence there is little consensus in the predictions. The input and output formats vary across the tools. It has been suggested that an ensemble approach that takes into account various prediction scores might perform better. There is a need for a tool that can run multiple such tools on a dataset in a more accessible and modular manner, whose output can then be combined to select consensus drivers.

We developed wrappers for various driver mutation predictions tools using Galaxy based framework. In order to perform predictions using multiple tools on the same dataset, we also developed Galaxy based workflows to convert VCF format to tool specific formats. The tools are publicly available at: https://github.com/saketkc/galaxy_tools The workflows are available at: https://github.com/saketkc/galaxy_tools/tree/master/workflows

## I. INTRODUCTION

Cancer is known to be arise from genomic aberrations[1]. With the advent of Next Generation Sequencing, it has been possible to profile the genomes from cancer patients and perform an in-depth analysis to yield insights that could be used for therapeutic, diagnostic and prognostic applications. Many of these profiles have also been released inn public domain, such as TCGA [2]. Given huge datasets such as these, the challenge has been to make sense out of them.

It has been suggested that Cancer is the outcome of Darwinian evolution at the cell level involving genetic variation. Natural selection acts on the phenotype level, thus positively selecting the cells which have acquired those mutations, that allow such cells to proliferate. Cells with such mutations lead to abnormal growth in the tumors. This growth may turn out to be controlled such as in the case of skin moles or it may end up leading to cancer tissues

These set of mutations are the ‘drivers’ of cancer that confer selective advantages to the cell to grow autonomously or alternatively to prevent cell death by affecting the apoptosis pathways ultimately leading to the positive selection of the cell. [3].

The ‘passengers’ do not confer any growth advantage to the cells.’Drivers’ thus, by definition are found in ‘cancer’ genes. the ‘passengers’ are known to be distributed randomly [3]. The ‘cancer genes’ are known to contribute

Identification of these set of driver mutations across various cancer types, would possibly act as a set of prognosis markers besides acting like therapeutic targets. Different approaches have been used to differentiate drivers from passengers. The challenge lies in correctly classifying thousands of mutations generated out of whole genome or exome sequencing as drivers or passengers.

Multiple approaches have been used for predicting driver mutations. These use information ranging from difference in mutational frequency among drivers and passengers to calculating functional impact scores , besides using a curated dataset of driver and passenger mutation for training classifiers [4].

Different tools use different methods and input formats. This often makes the task of biologists difficult. These tools, owing to different underlying approach used for prediction give non-consensus predictions for the same set of mutations. Galaxy [5, 6, 7] is an open source framework that allows running Bioinformatics tools in a reproducible manner. Galaxy thus provides a user friendly graphical interface accessible through a web browser for running such tools in a reproducible manner. Galaxy is used extensively for anlayzing next-generation sequencing datasets[8]. However few driver mutation predictor tools have been developed for Galaxy.

## II. METHODS

We developed wrappers around tools described in Table 1 leveraging Galaxy’s Toolshed framework[9]. These tools are freely available from the toolshed https://toolshed.g2.bx.psu.edu/ and are open source.

**Tabel 1:**
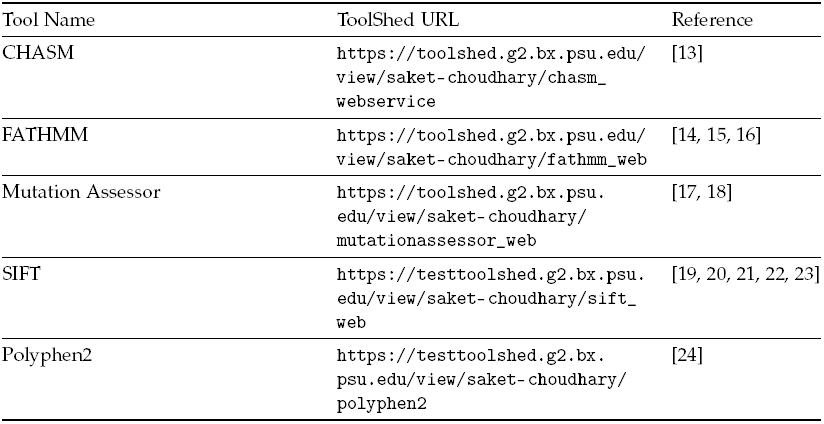
Tools Wrapped in Galaxy

One of the standard formats used to store mutation data is the VCF(Variant Call Format) [10]. In order to streamline the process of generating tool specific inputs, we developed workflows in Galaxy that take a VCF as an input and convert it to the tool specific input format.

There have been previous attempts at comparing such softwares [11] [4]. It has also been suggested that an aggregate approach such as the Condel[12] program which uses a weighted average score of SIFT, Polyphen and Mutation Assessor to make predictions and is shown to outperform each individual method. GalDrive aims at streamlining the process of comparing the methods by providing tools in Galaxy that can be used to run such comparison pipelines in a reproducible manner

## III. RESULTS & DISCUSSION

The GalDrive toolbox provides set of tools and workflows for running comparative pipelines for driver mutation prediction in a reproducible and flexible manner. Utilizing Galaxy’s set of internal tools with certain custom developed tools we created workflows that allows direct utilization of VCF file, which is a more standard format as inputs to these workflows. This can serve as a powerful tool to the biologists, who might not be acquainted with command line tools to run the methods, rest aside the need to perform various format inter-conversions.

These workflows can further be utilized to generate ensemble scores by combining output scores of two or more tools’ output.

## IV. ACKNOWLEDGMENTS

We would like to thank the Galaxy Team for their valuable help on the mailing-list during the process of development.

